# Predicting sites of epitranscriptome modifications using unsupervised representation learning based on generative adversarial networks

**DOI:** 10.1101/2020.04.28.067231

**Authors:** Sirajul Salekin, Milad Mostavi, Yu-Chiao Chiu, Yidong Chen, Jianqiu (Michelle) Zhang, Yufei Huang

## Abstract

Epitranscriptome is an exciting area that studies different types of modifications in transcripts and the prediction of such modification sites from the transcript sequence is of significant interest. However, the scarcity of positive sites for most modifications imposes critical challenges for training robust algorithms. To circumvent this problem, we propose MR-GAN, a generative adversarial network (GAN) based model, which is trained in an unsupervised fashion on the entire pre-mRNA sequences to learn a low dimensional embedding of transcriptomic sequences. MR-GAN was then applied to extract embeddings of the sequences in a training dataset we created for eight epitranscriptome modifications, including m^6^A, m^1^A, m^1^G, m^2^G, m^5^C, m^5^U, 2′-*O*-Me, Pseudouridine (Ψ) and Dihydrouridine (D), of which the positive samples are very limited. Prediction models were trained based on the embeddings extracted by MR-GAN. We compared the prediction performance with the one-hot encoding of the training sequences and SRAMP, a state-of-the-art m^6^A site prediction algorithm and demonstrated that the learned embeddings outperform one-hot encoding by a significant margin for up to 15% improvement. Using MR-GAN, we also investigated the sequence motifs for each modification type and uncovered known motifs as well as new motifs not possible with sequences directly. The results demonstrated that transcriptome features extracted using unsupervised learning could lead to high precision for predicting multiple types of epitranscriptome modifications, even when the data size is small and extremely imbalanced.

## INTRODUCTION

Epitranscriptome is an exciting emerging area that studies modifications in transcripts. The insurgent interest is largely fuelled by the recent discovery of widespread N6-methyl-adenosine (m^6^A) methylation in mammalian mRNAs [1, 2], which has been shown to play important regulatory roles in every stage of RNA metabolism and involve in many diseases. Besides m^6^A, many other types of modifications are found to exist in the eukaryotic transcriptome. While some of them including N1-methyladenosine (m^1^A), 5-hydroxymethylcytosine (hm^5^C), 5-methylcytidine (m^5^C), 2′-O-methylation (2’-O) and pseudouridine (Ψ) are found widespread, other types such as Dihydrouridine (D), and m^2^G have only a hundred sites discovered thus far. These exciting findings have spurred intense research to identify transcriptome modifications in different cells and to decipher their roles in regulating various biological processes [3].

We consider in this paper the prediction of transcriptome modification sites from transcript sequences. This problem is naturally a supervised learning task, which aims to train predictive models for each type by using labeled positive and negative modification sites. There is a large collection of algorithms for predicting m^6^A sites from mRNA sequences[4–10], most notably SRAMP. However, such predictive algorithms for other modifications are still scarce because training robust models for these modification sites face several challenges[11, 12]. First, training for modification types with scarce labeled samples suffers from significant overfitting, making the model incapable of learning true methylation-specific sequence features from random noise patterns. Second, training data for all modifications suffer the significant class imbalance, a common challenge in most of the genomics applications, where only a small percentage of transcriptome nucleotides are true methylation sites and the majority of them are negatively labeled unmodified sites. Unfortunately, traditional supervised learning tends to treat these extremely small numbers of instances as noise and again fails to learn desired methylation-specific sequence features from the positive samples[13]. Third, predicting modification sites of these different types altogether imposes a greater challenge as different types are likely to share similar biological sequence patterns making them less distinguishable from each other. One viable solution to address these challenges is to take advantage of the vast unlabelled part of the transcriptome sequences with unsupervised representation learning to learn transcriptome-wide sequence features as a whole. We assume that the unlabelled part of the transcriptome could contain unidentified modification sites and/or modification related functional sites (e.g., RNA binding protein binding sites). Therefore, the unsupervised learning of transcriptome sequences may help capture modification-related features, which are difficult to learn otherwise by a supervised approach using only labeled sequences. We intend to learn these features in an unsupervised setting to leverage the supervised learning using labeled data such that the classifiers trained using only a few labeled examples can generalize to predict modification sites robustly. Unsupervised learning methods have recently attracted an influx of research interest, especially in the field of bioinformatics. For example, Restricted Boltzmann machines have been applied for unsupervised pre-training of neural networks that were later used to initialize the supervised learning of protein 3D structures [14, 15] and amino acid contacts[16]. Besides, Asgari et al. [17] proposed a low dimension representation method for protein sequences, which is inspired by the widely known word2vec model of natural language processing. However, none of these methods have been trained to extract features from RNA sequences.

In pursuit of addressing the challenges described above, we propose an unsupervised feature construction approach based on Generative Adversarial Network (GAN) [18]. Our method, aptly known as MR-GAN, predicts multiple types of <u>m</u>odification sites in <u>R</u>NA using <u>GAN</u> based unsupervised feature learning. This method delves deep into the largely unexplored area of low dimension representation of RNA sequences and demonstrates the usefulness of the unsupervised feature learning in handling some of the most difficult problems in the intersection of machine learning and bioinformatics (small and imbalance dataset). Though the idea of GAN has been fervently researched in the computer vision domain in recent years [19–21], this architecture is largely uncharted territory for bioinformatics except a few [22, 23]. The original GAN framework [18], which consists of two modules known as generator and discriminator, was designed to learn a generative distribution of data through a two-player minimax game. While the generator’s goal is to “fool” the discriminator by producing samples that are as close to the real data distribution as possible, the discriminator strives to be not fooled by correctly classifying between real and fake data. The adversarial framework has been shown to provide a superior loss to other traditional ones based on mean square error or mutual information for learning data distributions.

Despite the success of GAN in mimicking the underlying data distribution, GANs cannot be readily applied to learn the abstract representation of data due to the lack of an efficient inference module. Hence, in order to learn the low dimension feature representation of RNA sequences using GAN based framework, we employed Adversarial Learned Inference (ALI) model [24], which jointly learns an encoder network and a decoder network using the adversarial process similar to GAN. The decoder network takes random noise as input and maps it to the data space, whereas the encoder network maps RNA sequences to the latent representation. Finally, the encoder, decoder, and a discriminator network get combined into the adversarial game, where the discriminator network is trained to distinguish between the joint distribution of latent/data-space samples from the decoder and encoder network.

We adapted the inference learning approach of ALI and employed it to a large compendium of the pre-mRNA sequence of the human transcriptome. The objective was to learn an abstract representation of transcriptomic sequences that are typically 51 bp long and utilize the embedding as feature vectors for predicting eight different types of transcriptome modification sites including m^1^A, m^1^G, m^2^G, m^5^C, 2’O, m^5^U, pseudouridine (Ψ), and Dihydrouridine (D). Evaluating the effectiveness of features by predicting multiple post-transcriptional modifications serves in the manifold. First, the computational prediction of distinct epitranscriptomic marks is biologically significant and a long-sought goal for bioinformatics researchers. Accurate identification of these marks is essential for deciphering their biological functions and mechanism. However, discriminating between RNA modifications using only genomic sequence is a challenging task because there are more than 100 different types of RNA modifications characterized so far in diverse RNA molecules, including mRNAs, tRNAs, rRNAs and lncRNAs and they may share similar nucleotide sequence preference. Many transcriptome-wide sequencing technologies have been developed recently to determine the global landscape of RNA modifications (e.g, Pseudo-seq, Ψ-seq, CeU-seq, Aza-IP, MeRIP-seq, m6A-seq, miCLIP, m6A-CLIP, RiboMeth-seq, Nm-seq and m1A-seq) that identifies distinct epitranscriptomic marks [25]. However, individual experimental identification of these modification sites is very costly, labor-intensive and time-consuming. So, we created a benchmark dataset by combining eight different types of post-transcriptional modification data (e.g., m^1^A, m^1^G, m^2^G, m^5^C, m^5^U, 2′-*O*, Ψ and D). We compared the effectiveness of the embeddings with the one-hot encoding of RNA sequences and demonstrated that the learned embeddings outperform one-hot encoding by a significant margin (Fig. 2). We have also applied our method on the m^6^A dataset provided by SRAMP and improved the performance by 4%-12% for different case scenarios in predicting m^6^A site (Fig. 3). Finally, we carried out exploratory analysis via t-SNE to rationalize the superiority of MR-GAN features as well as investigated motifs learned by our model to determine the biological relevance of the embedded representation.

## MATERIAL AND METHODS

### Transcriptome dataset for training MR-GAN

Because our goal is to learn a low dimensional representation of transcriptome sequences using GAN, we employed pre-mRNA sequences from the entire human transcriptome to train the proposed MR-GAN model. Engaging the large corpus of pre-mRNA sequences is important as it aids our unsupervised training to learn patterns from coding and noncoding region to ensure that sufficient contexts are observed. The training dataset was compiled from all the chromosomes of the human genome hg38 assembly [26]. Separate fasta format files (*.fa) were downloaded from the UCSC genome browser for each of the chromosomes. Each of the files comprises of intronic and exonic sequences for all the genes belonging to the specific chromosome. We then chopped the sequences at the non-overlapping interval of every 51 bp and got rid of the sequences that contain character N (represents ambiguous nucleotide). We selected 51 bp as the input RNA sample length because previous studies involving the prediction of RNA modifications have discovered this length as preferable for capturing contextual information [5, 6]. This preprocessing step results in a total of 40.7 million samples of length 51 bp. Finally, in order to feed into the MR-GAN for unsupervised learning, each of these sequences was represented by a 4 × 51 binary one-hot encoded matrix with rows corresponding to A, C, G, and U.

### Datasets for epitranscriptome modifications

To train and evaluate models for human transcriptome modification site prediction using transcript sequence features extracted by the MR-GAN encoder, we created two benchmark datasets. The first dataset includes sites from eight different types of transcriptome modification and negative random sequences (see Table 1). The positive sites were collected primarily from RMBase [27], which provides location information of the modified single base. We utilized the modified single base as the center and extended it to 51 bp by including 25 bp nucleotide sequences from both the upstream and downstream direction to form positive sequences. We also extracted a set of background sequence samples, randomly sampled from the genomic locations that do not contain any modification sites; we ensured that the proportions of the sequences centered at either A, U, C or G are roughly balanced (see Table 1). Then, the negative training samples were created for different types of modifications separately. For 2’-O or Pseudouridine, which occur on all four different nucleotides, the negative samples include the background samples and samples from all other eight types of modification. Otherwise, for any of the other seven modifications that are associated with a unique nucleotide, the negative samples include only sequences centered at that unique nucleotide from the background samples and those from other types. While some of the modifications have large positive sample sizes, the majority of others as in m^2^G and m^1^G have very few positive samples. Training reliable prediction models with small, highly imbalanced training datasets are highly challenging leaning tasks. In this work, we show the power of the MR-GAN encoder in helping extract discriminate sequencing features for different modification sites.

**Table 1:** Number of samples for different transcriptome modifications in the benchmark dataset.

| Modification type | Number of samples | Center nucleotide |  |  |  |
| --- | --- | --- | --- | --- | --- |
|  |  | A | C | G | U |
| 2'-O | 2802 | 422 | 820 | 899 | 661 |
| D | 162 |  |  |  | 162 |
| m<sup>1</sup>A | 3173 | 3173 |  |  |  |
| m<sup>1</sup>G | 29 |  |  | 29 |  |
| m<sup>2</sup>G | 59 |  |  | 59 |  |
| m<sup>5</sup>C | 536 |  | 536 |  |  |
| m<sup>5</sup>U | 30 |  |  |  | 30 |
| Ψ | 3732 | 31 | 40 | 29 | 3632 |
| Negative (random) | 7699 | 1963 | 1815 | 1917 | 2004 |
| Total | 18222 | 5589 | 3211 | 2933 | 6489 |

The second dataset consists of positive and negative samples for transcriptome m^6^A methylation. We choose to train an MR-GAN on m^6^A separately because m^6^A is the most abundant and widely studied transcriptome modification and there are also existing sequence-based prediction algorithms. This dataset was derived mostly from [6], which used a similar dataset to train an m^6^A site predictor called SRAMP. Positive samples of this dataset are miCLIP sites from [28] that also contain the m^6^A motif (DRACH) and the negative samples are random sequences that also have the DRACH motif located at the center but have no miCLIP detected m^6^A sites. Similar to for SRAMP, two sets of training data, namely the full transcript mode, where the training sequences were collected from the full transcripts, and the mature mRNA mode, which extracted training sequences from cDNA sequences, were prepared. Table 2 provides detailed information about the m^6^A training datasets.

**Table 2:** Numbers of samples in the training and testing dataset for m^6^A. Data were prepared for the full transcript and mature mRNA modes.

|  | Full transcript mode | Mature mRNA mode |
| --- | --- | --- |
| Training data (pos:neg) | 31901:229204 | 26755:51094 |
| Testing data (pos:neg) | 8284:31070 | 6905:34349 |

### Generative adversarial network

GAN, proposed by Goodfellow et al. [18], is a generative network that learns the distribution of data and produces samples of synthesized data from the captured distribution. GAN includes two differentiable functions characterized by neural networks: the discriminator function *C*(*x*; *θ*_*C*_)with parameters *θ*_*C*_ that outputs a single scalar representing the probability of *x*. from the real rather than synthesized data distribution and the generator *D*(*z*; *θ_D_*) parameterized by *θ*_*D*_ that maps samples from a prior of input noise variables *Pz*(*z*) to data distribution *p*_data_(*x*). GAN is trained by implementing a two-player minimax game, where *D* is optimized to tell apart real from synthesized or fake data and *D* is optimized to generate data (from noise) that “fools” the discriminator. This joint optimization can be formulated as

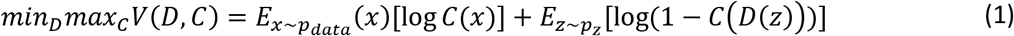

Despite the recent successes of GAN in computer vision, the idea was unexplored in the genomics domain until Frey et al. [22] applied the framework on DNA sequences with some modifications. Instead of training a generative model that produces realistic DNA sequences, they synthesize sequences with certain desired properties. For instance, the data distribution captured by GAN was utilized to design DNA sequences with higher protein binding affinity than those real sequences found in the protein binding microarray (PBM) data.

### MR-GAN for unsupervised learning of transcriptome sequences

As discussed earlier, GANs lack an inference network that prohibits them from understanding abstract data representations. People have used the discriminator network to extract features but as a learning entity to separate the real and synthetic samples, the discriminator primarily learns discriminate features between real and synthetic samples. Hence, the discriminator features are not a true representation of the underlying data. Learning an inverse mapping from generated data *E*(*x*) back to the latent input *z* can be one viable solution to the problem under consideration. To this end, we propose MR-GAN, a model inspired by the ALI and BiGAN framework, which consists of three multilayer neural networks as depicted in Figure 1. Briefly, in addition to the generator *D* (or decoder in this case) from the standard GAN framework, MR-GAN includes an encoder *E*, which maps data *x* to latent representations *z* . The discriminator *C* of MR-GAN discriminates joint samples of the data and corresponding latent component (pairs (*x*; *E*(*x*)) versus (*D*(*z*); *z*) as real fake pairs, where the latent variable is either an encoder output *E*(*x*) or a generator input *z*. Overall, the training is performed to optimize the following minimax objective.

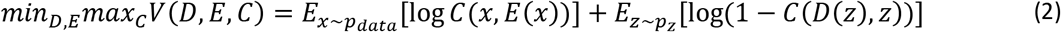

**Figure 1:**
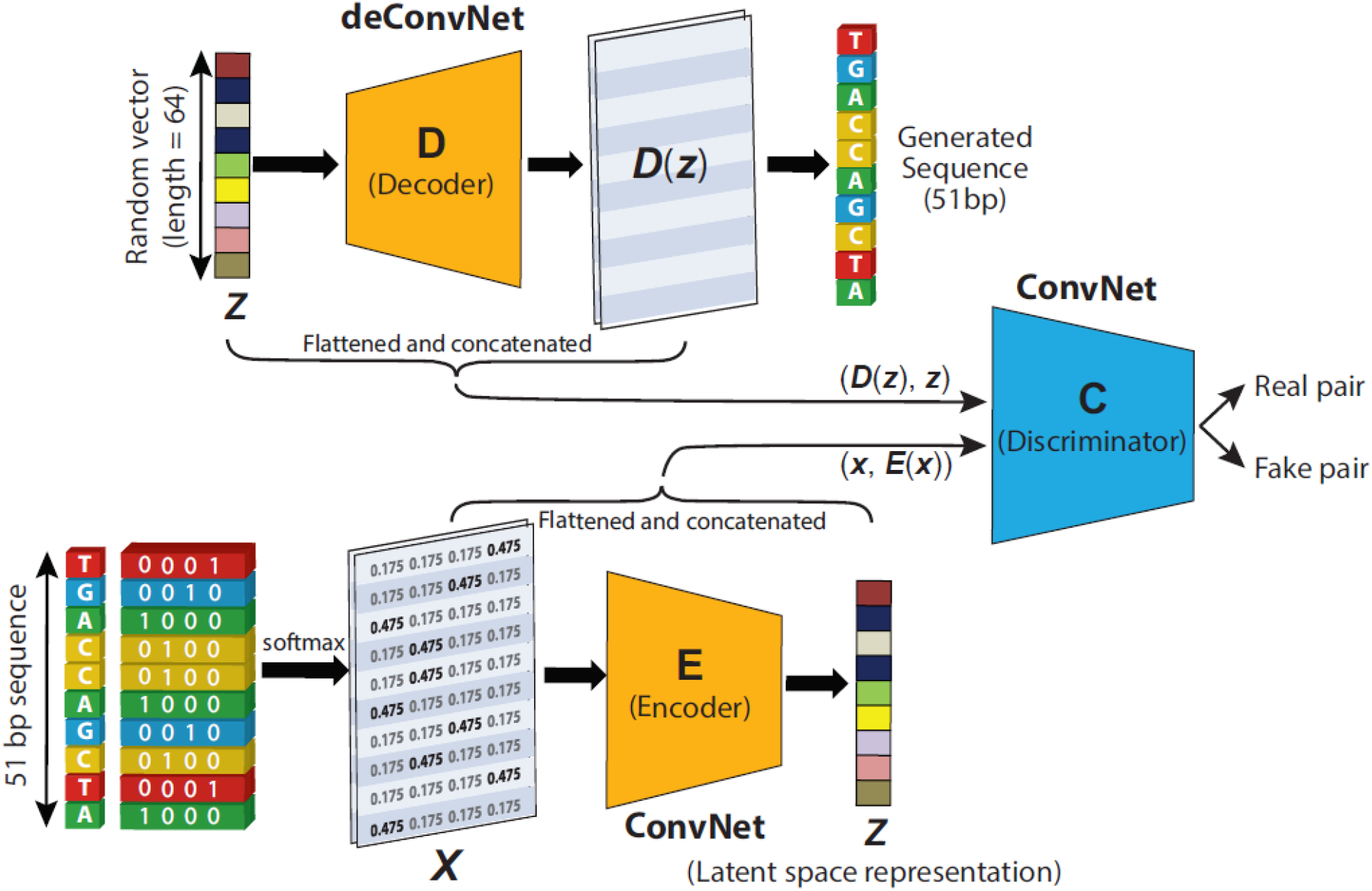
Unsupervised Feature Learning Scheme of MR-GAN

In MR-GAN’s setting, aside from training a generator, we train an encoder *Q*: Ω(*x*) → Ω(*z*), which maps data points *x* into the latent feature space. The discriminator takes input from both the data and latent representation, producing *P*_*C*_ (*Y*|*x*, *z*), where *Y* = 1 if *x* is sampled from the real data distribution *p*_*x*_, and *Y* = 0 if *x* is generated through the output of *D*(*z*), and *z* ∼ *p*_*z*_ . Each of the *C*, *D*, and *E* modules, which are parametric functions, are optimized simultaneously using a stochastic gradient descent algorithm. The specific architecture of *C*, *D*, and *E* modules can be found in supplementary Table 1. Since the MR-GAN encoder learns to capture the semantic attributes of whole human transcriptome sequences, it produces more powerful feature representations than a fully supervised model that is trained using only the labeled transcriptome modification site sequences, especially for those occasions where training data are limited.

### Training MR-GAN

The GAN framework trains a generator, such that no discriminative model can distinguish samples of the data distribution from samples of the generative distribution. Both generator and discriminator are trained using the optimization function noted in equation (1). In the MR-GAN setting, the loss function (2) is considered, which is optimized over the generator, encoder, and discriminator. However, both (1) and (2) can be considered as computing the Jensen-Shannon divergence, which is potentially not continuous concerning the generator’s parameters, thus leading to poor training convergence and stability [29]. In [29], an alternative optimization function was proposed that is based on the Earth-Mover (also called Wasserstein-1) distance *W*(*q*, *p*)and is defined as the minimum cost of transporting mass in order to transform the distribution *q* into the distribution*p.*In contrast to equation (1) and (2), the Wasserstein distance is a continuous and differentiable function in the parameter space and therefore has improved training stability. However, it requires weight clipping to be implemented that still leads to the generation of poor samples and failure to converge. Therefore, we trained MR-GAN using Wasserstein optimization function but instead of enforcing the weight clipping, we imposed a soft version of the constraint with a penalty on the gradient norm as follows

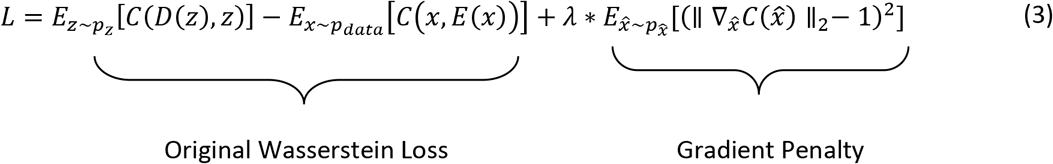

This technique is known as WGAN-GP (Gradient Penalty) and was initially proposed in [30]. The entire MR-GAN framework was optimized using the standard stochastic gradient descent method, where the learning rate was set to 1e-4. In our implementation, the batch size was equal to 100 samples. We trained the network for two epochs, which takes around four days in a computer with a single GPU. In our experience, the loss keeps oscillating during the training process as has been observed in previous studies [31–33]. However, WGAN-GP has shown that the discriminator loss is close to zero when the GAN model approaches convergence. As a result, we took a snapshot of our model at every 2,000 iterations during the training process and selected the model with minimum discriminator loss (−5.2) for downstream processing. In our training, the minimum discriminator loss occurs at iteration 809,600, whereas the total number of iterations for two epochs was 814,600.

### Training predictors for epitranscriptome sites based on features extracted from the MR-GAN encoder

After the MR-GAN model is trained in an unsupervised manner using transcriptome-wide pre-mRNA sequences, we retained the MR-GAN encoder and used it as a feature extractor for modification site sequences and trained prediction models based on these features [21]. To train a model to predict modification sites, we first fed each 51 bp long sequences in our training and testing datasets into the encoder and concatenated the feature maps from the last four convolutional layers into a vector of 2,112 features. Then, support vector machine (SVM) classifiers with the Radial basis function kernel were trained as the site predictors for each modification type based on the feature vectors extracted from the corresponding training sets. This resulted in eight predictors for the first dataset and a separate predictor for m^6^A. Note that these eight predictors together can also be considered as a multi-class classifier trained based on the one-vs-rest approach.

To obtain a baseline prediction performance, we also trained SVM classifiers using the one-hot encoded modification sequences. One hot encoding is a widely used encoding approach in deep learning to represent biological sequences with numeric formats [34–37]. The one-hot encoding converts a 51 bp mRNA sequence into a 4×51 binary matrix, where each row corresponds to either A, C, G or U and a single “1” in each column encodes the corresponding nucleotide at that location of the sequence.

## RESULTS

### Prediction performance of the eight epitranscriptome modifications

In this section, we comprehensively evaluate the performances of the eight modification predictors trained on MR-GAN generated features and the one-hot encoding using the first dataset. Because the positive and negative samples for any modification are heavily imbalanced, where negative samples are about 3 to 200 times more than the positive samples, the appropriate metric to gauge the classification performance is the area under the precision-recall curve (auPRC). Since both precision and recall are dependent on the number of true positives rather than true negatives, the auPRC is less prone to inflation by the class imbalance than auROC [38]. Fig. 2 shows the auPRC performances achieved by a 5-fold cross-validation scheme for each of the modifications. As described earlier, each modification predictor was trained by selecting the samples of the particular modification as the positive class whereas the samples having the same center nucleotide as a positive class from the rest of the modifications including the random negative ones were attributed to the negative class. It is clear from Fig. 2 that MR-GAN extracted features outperform one-hot encoding representation by significant margins for most of the RNA modifications. The auPRC increase attained by MR-GAN features ranges from 5.6% to 19.2% across different modifications. Notably, the performance improvements are more prominent for the modifications with a comparatively lower number of samples. For instance, the average auPRC for predicting D, m^2^G, m^5^C and m^5^U using MR-GAN embedding is 12.3% higher than the one-hot encoding comparing to the 3.8% average auPRC increment for the rest of the modifications with larger sample size (2’-O-Me, Pseudouridine, m1A). This indicates that the unsupervised training of MR-GAN has helped the model to learn additional information that otherwise would be difficult to learn with a limited number of training samples. Furthermore, the MR-GAN features consistently deliver prediction results at higher than 85% auPRC which is a remarkable feat considering the minuscule ratio of positive to negative samples in the heavily imbalanced dataset. Using a highly skewed dataset to train predictors generally leads to the outcome that most of the positive samples are misclassified as negative ones. To solve this complexity, many prediction algorithms resample the negative data to balance the ratio of positive and negative subsets. Remarkably, we did not perform any data balancing technique in training, as the features learned by the MR-GAN encoder were powerful enough to handle the class imbalance problem by itself. Finally, we noticed that the one-hot encoded feature representation performs comparably to the MR-GAN features for two modification types (m^1^A and m^1^G). Because there are only 29 positive samples for m^1^G, it is reasonable to expect that the one-hot encoded feature representation and the MR-GAN features reach similar performance but these performances might not be generalized due to the small sample size. For m^1^A, where there are 3,173 positive training samples, it is likely that the positional sequence pattern in training samples already contains significant discriminating information for the one-hot predictor to reach maximum accuracy. Taken together, these results demonstrate that the proposed unsupervised representation by MR-GAN successfully captures the important features required for discriminating the RNA modifications, which might not be captured with small training samples.

**Figure 2.**
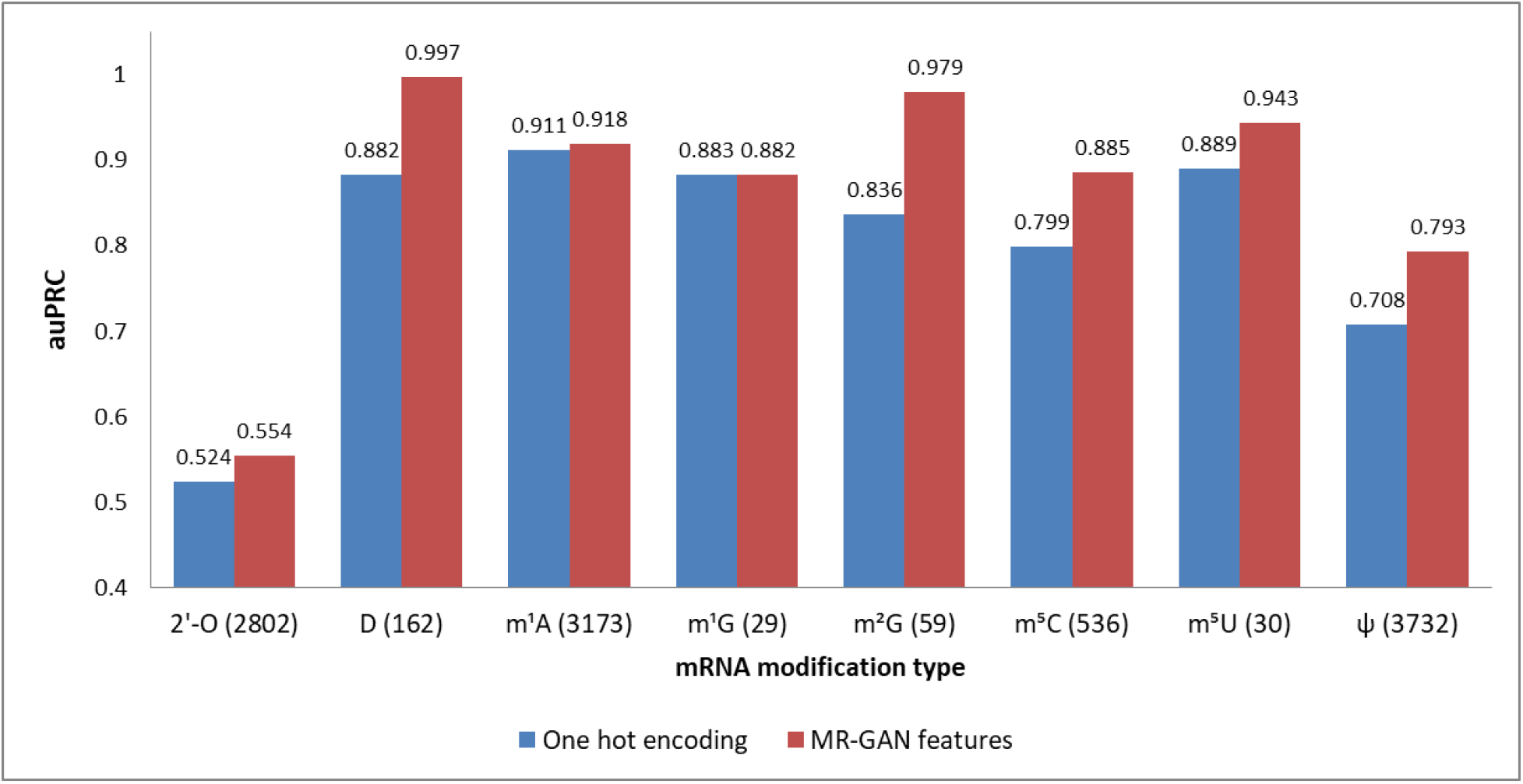
Prediction performance (auPRC) of eight epitranscriptome modifications using MR-GAN features and one-hot encoding. The number next to a modification name is the number of corresponding positive samples. The auPRCs are listed on top of the bars.

### Prediction performance of m^6^A sites

While in the previous section we demonstrated the superior prediction performance of MR-GAN embedding on modifications with relatively smaller training samples, we investigate in this section whether this unsupervised feature learning can help improve the performance of m^6^A prediction, where the number of positive samples required for training is large. Because of the availability of m^6^A data from SRMAP, we were able to evaluate the performance on independent testing datasets as opposed to cross-validation. Similar to the SRAMP evaluations in [6], we investigated the MR-GAN model performance in terms of auPRC on the full transcript mode and mature RNA mode using the independent testing data as described in Table 2.

Figure 3 summarizes the prediction results of MR-GAN and SRAMP on testing data for both the input modes. As evident, the MR-GAN extracted features outperform SRAMP proposed encodings in both modes (11.1% for the mature mRNA mode and 3.6% for the full transcript mode), suggesting that unsupervised learning from the entire transcriptome sequence can improve the learning of modification-related features over the encodings employed in SRAMP algorithm. It is pointed out in [6] that SRAMP suffers in the mature mRNA mode due to the discarding of all introns, which may disrupt the original sequence context of an m^6^A site and therefore reduce the discriminative capability of the extracted features. Also, the distance between an m^6^A site and a non-m^6^A site generally becomes closer in mature mRNA sequences compared with that in the corresponding pre-mRNA sequences, which further aggravates the prediction outcome of SRAMP. By contrast, the unsupervised training enables MR-GAN to learn the information about pre-mRNA sequences and thus produce considerable prediction improvement.

**Figure 3.**
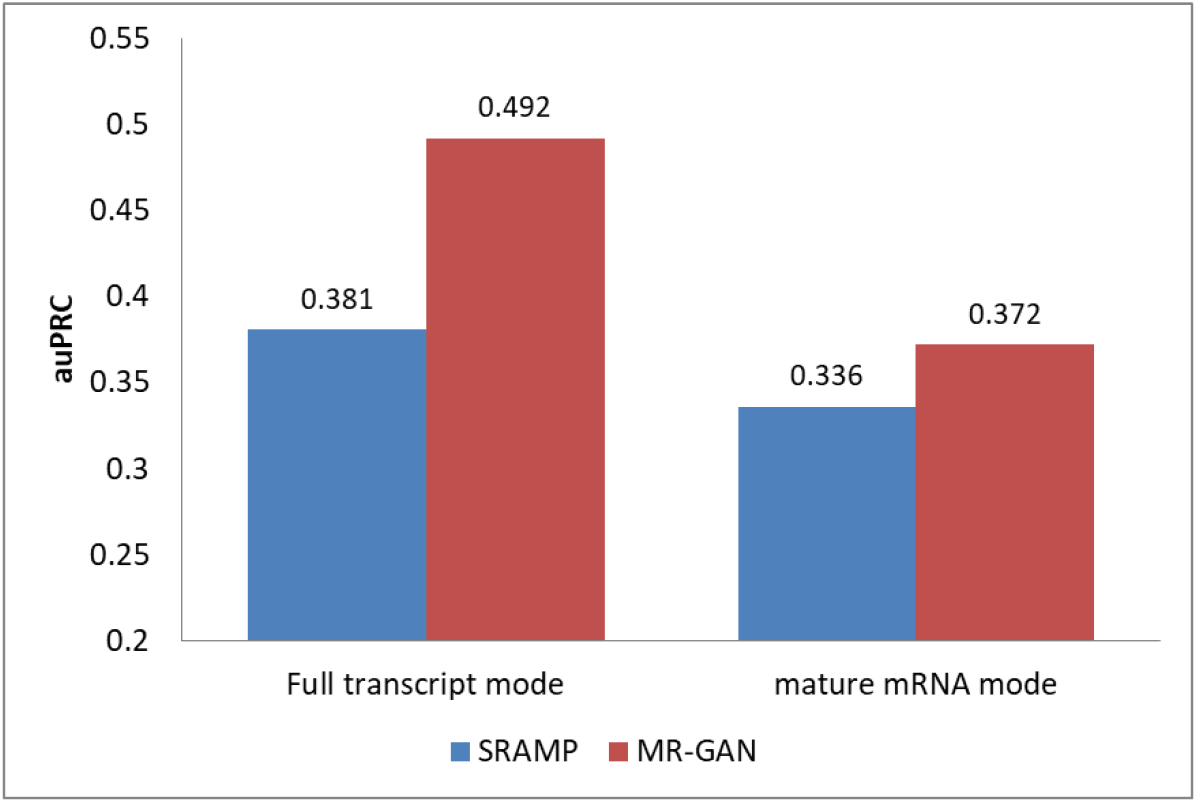
m^6^A prediction performance (auPRCs) of MR-GAN and SRAMP

### Visualizing MR-GAN features for different epitranscriptome modifications

To gain an intuitive understanding of the superior performance of MR-GAN features and explore the relationship among features of different modifications, we applied t-SNE [39] to project the high dimensional features onto 2D and 3D space. t-SNE is a popular nonlinear dimension reduction method that optimizes the similarities of the probabilities of distances between high dimensional samples space and corresponding lower-dimensional projected samples. It has been widely used for data visualization in a variety of fields such as video, image, and audio signals [39]. For this visualization, we first utilized principal component analysis (PCA) to map the MR-GAN and one-hot encoded representations of different RNA modification samples to lower dimensions, which were then fed to t-SNE to obtain representations in 2D and 3D spaces. We first compared the t-SNE plots in a 3D space between MR-GAN features and the one-hot encodings (Fig. 4). For this comparison, we only included the five epitranscriptome modifications with a comparatively larger number of samples. As evident in Fig. 4(a), all one-hot encoded samples were squeezed together and barely separable. In contrast, MR-GAN extracted features show well-separated groups of samples (Fig. 4(b)). As a result, it is much easier to achieve separable hyperplanes in the SVM classifier when MR-GAN features are used.

**Figure 4.**
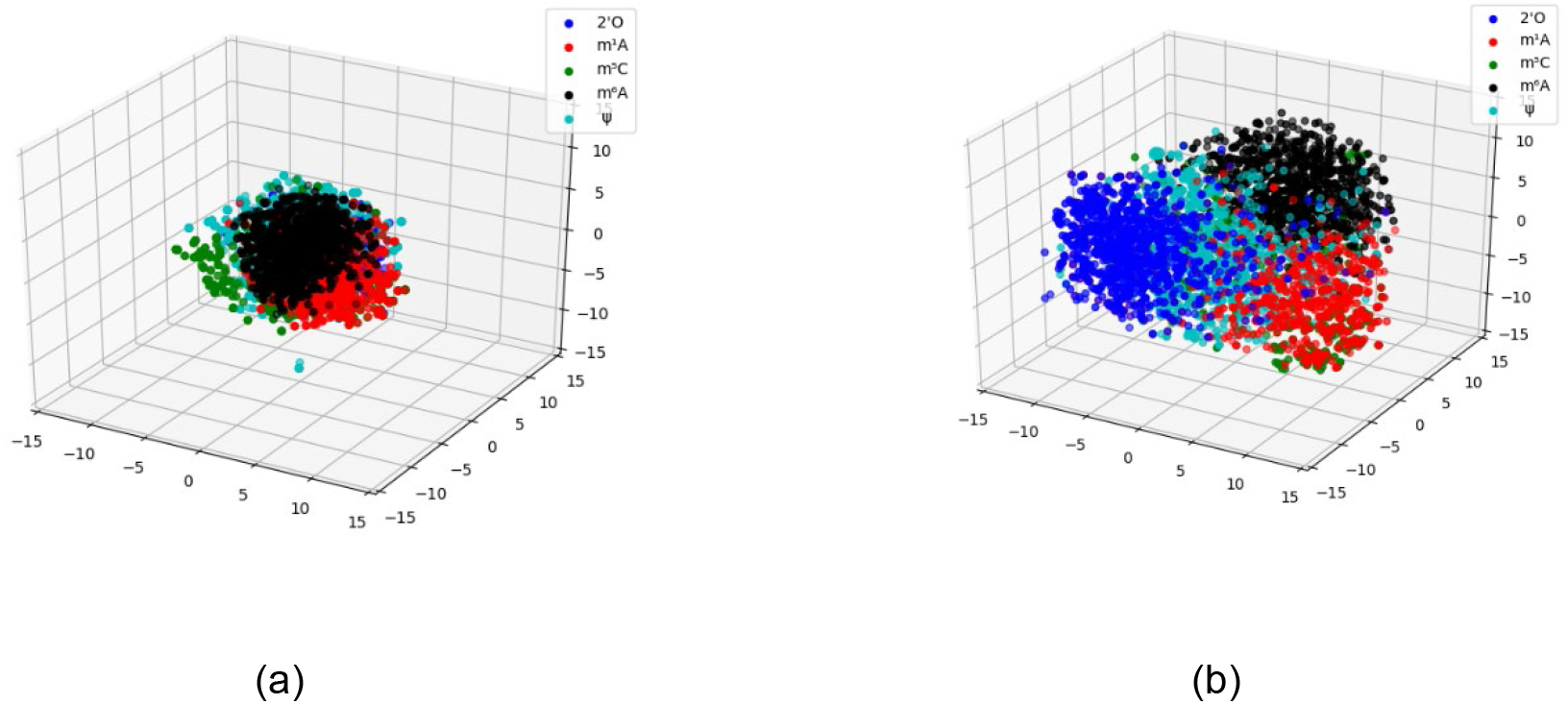
3D scatterplots of t-SNE projected (a) one-hot encoded samples and (b) MR-GAN features for five epitranscriptome modifications that have a comparatively larger number of samples.

One way to validate the data embedding is to verify if the samples with similar functionalities are clustered together in the embedding space. Therefore, we investigated the relationship of MR-GAN features between different modifications, where we projected the data samples of eight types of modifications to the 2D space using t-SNE (Fig. 5). To produce a clear and uncluttered 2D plot, we randomly picked a maximum of 70 samples from each of the modifications. Fig. 5 reveals that the four most abundant mRNA modifications, m^6^A, m^1^A, m^5^C, and 2’-O, each form unique groupings with somewhat small overlapping among them, suggesting that each of them has its distinct sequence patterns that might be associated with modification-specific RNA binding proteins and hence unique functions. Such distinct grouping is supported by their different transcript distributions, where m^6^A, m^1^A, m^5^C, and 2’-O are mostly enriched in the stop codon, the start codon, the 3’UTR and the CDS, respectively. However, there is a small overlapping between m^6^A and 2’-O. Indeed, single-base mapping technology has found that the second nucleotide from the 5’ cap of certain mRNA has *N*^6^, 2’-O-dimethyladenosine (m^6^Am) [28]. Moreover, 2’-O exists in all four types of nucleotides and it is not surprising to see 2’-O to share some overlapping with almost all other modifications. We also observe that the m^1^A cluster contains virtually all m^1^G samples, implying that m^1^A and m^1^G have significantly share sequence patterns and potentially similar regulatory functions. Certainly, m^1^A is shown to disrupt the Watson-Crick base-pairing and found to collaborate with m^1^G to induce local duplex melting in RNA [40]. The fact that they are isolated from other modifications may suggest that disrupting the base-pairing might be their unique function. Compared with the other three clusters, the m^5^C cluster shows the most overlapping with m^6^A and 2’-O. Although the evidence of collaborations between these three methylations is limited, m^6^A and m^5^C have been shown to enhance the translation of p21 by jointly methylating its 3’UTR. In addition to these four clusters, Ψ shows overlapping with m^6^A, m^5^C, and 2’-O. Ψ is found throughout different regions in mRNA. The samples of the remaining two modifications, D and m^2^G, are widely scattered without any clear patterns. They are also much less studied and their actual distributions in mRNA and their functions are mostly unknown.

**Figure 5.**
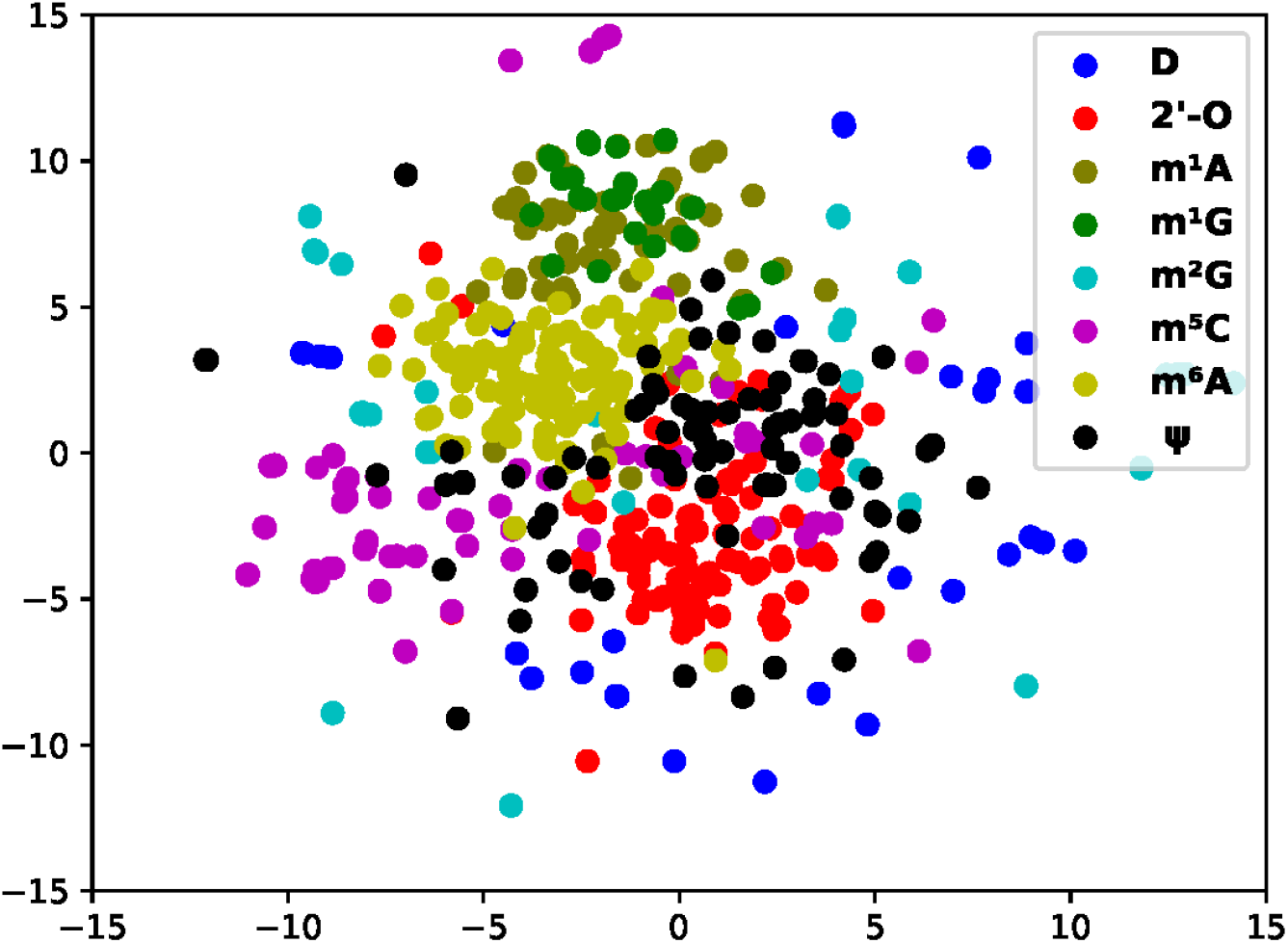
2D scatter plot of all modification sites using MR-GAN features map modifications with similar functions altogether

### Features learned by MR-GAN confirmed known modification sequence motifs

In this section, we delve into the encoder network to comprehend the sequence motifs for different mRNA modifications captured by the latent semantic representation of MR-GAN. Following a similar strategy described in DeepBind [34] for the sequence logo generation, we extracted the subsequences that give the maximum response in the convolutional operation for a kernel at the first layer of the encoder network. We repeated this subsequence extraction task on each of the sequences of the benchmark dataset using the 32 kernels at the first layer. Next, we performed motif enrichment analysis using MEME-ChIP [41] by feeding the unique subsequences from each modification type as the positive set while the subsequences extracted from the random samples of the benchmark dataset were utilized as the control set for each of the modifications to ensure the common background. The top-3 enriched motifs discovered by MEME-ChIP and the associated RNA binding proteins (RBPs) for the relatively abundant mRNA modifications are shown in Fig. 6. Two of the most enriched motifs detected for m^6^A belong to the well-known DRACH motif. The RBPs that share the motifs include FMR1, whose binding mRNAs in the mouse brain have been shown to be significantly methylated with m^6^A marks [42]. Other RBPs associated with these top motifs include splicing factors SRSF1 and Zinc finger CCCH domain-containing protein ZC3H10. Currently, no evidence of their interaction with m^6^A exist. However, m^6^A reader protein YTHDC1 is shown to regulate splicing through interacting with splicing factor SRSF3 and SRSF10 [43, 44] and the Zinc finger CCCH domain-containing protein ZC3H13 is reported to regulate m^6^A to control embryonic stem cell self-renewal. These existing results may point to unknown interactions of m^6^A with SRSF1 and ZC3H10. Next, we examined the known motifs for each of the remaining abundant modifications in the top enriched motifs and reported many other new motifs associated with known RBPs. To systematically investigate the associated RPBs predicted by MR-GAN, we accumulated the list of RBPs for each of the mRNA modifications that were identified as discriminative by MEME-ChIP in the above experiment, resulting in total 72 RBPs. We then performed a clustergram analysis (Fig. 7) such that the RBPs are clustered according to the significance with which they are found to be unique to these mRNA modifications by MEME-ChIP. Interestingly, we discovered that most of the modifications have at least one unique RBPs (Supplementary Table 2). For instance, RBM3 and RALY are unique to m^6^A, HNRNPC is unique to m^1^A, and SRSF2 is unique to m^5^C. RBM3 is shown to be a functional partner of the splicing factor SRSF3 [45], which is recruited by m^6^A to regulate alternative splicing [44]. RALY is a member of the hnRNP family, which are considered as indirect m^6^A readers [46, 47]. Also, there were also RBPs, such as SNRNP70, that are determined as discriminative for most of the modifications. SNRNP70 is a key early regulator of 5’ splice site selection. This result could suggest that regulating splicing would be a common function which various modifications possess. We also discovered that 55 of the 72 RBPs identified by this analysis overlaps with the proteins isolated by the RICK experiment [48], which systematically captures proteins bound to a wide range of RNAs (Supplementary Table 3). This indicates that our unsupervised learning captures RBPs that are biologically meaningful and repeatedly identified by other related studies.

**Figure 6.**
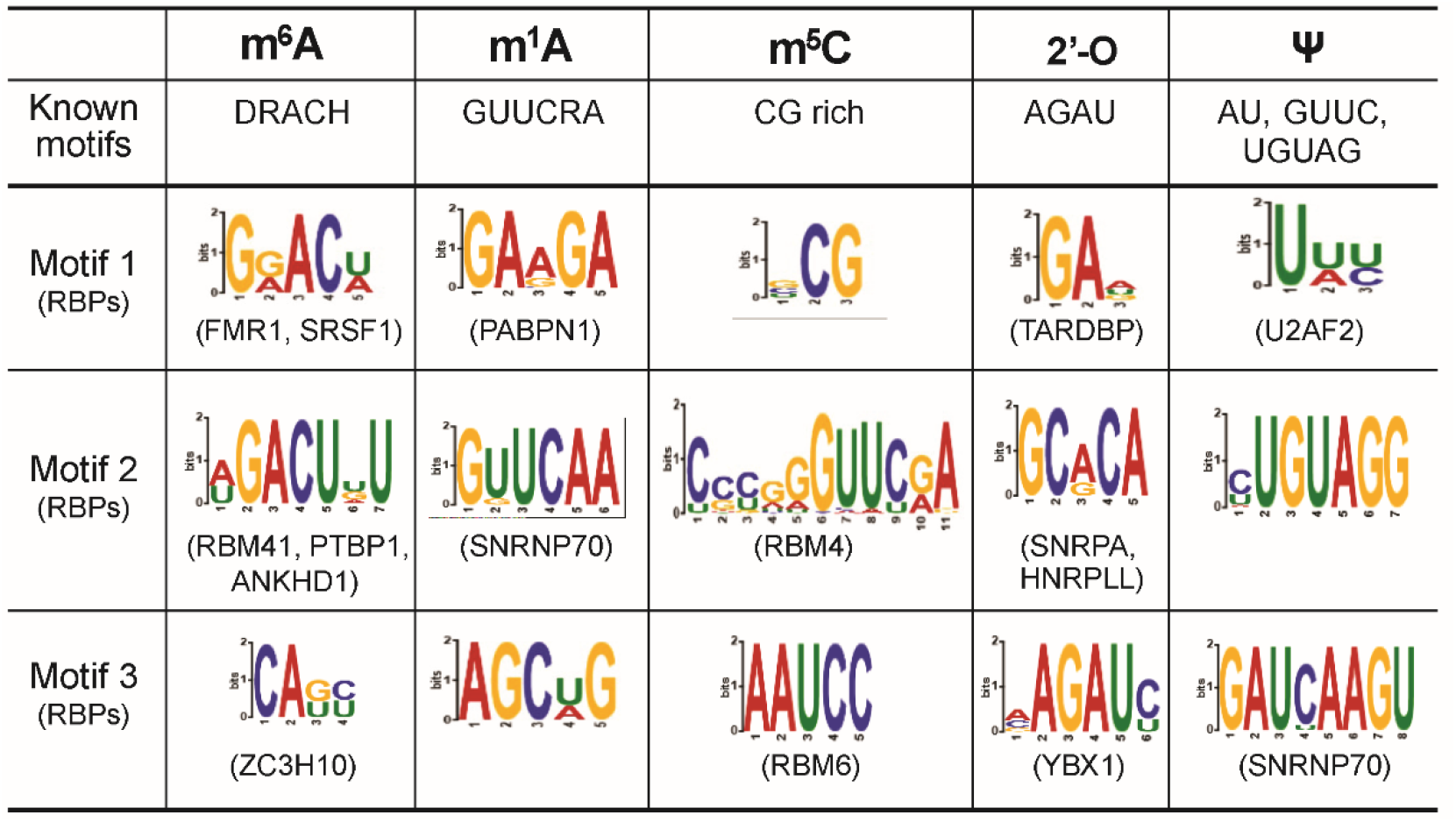
The top motifs of different transcriptome modifications learned by MR-GAN

**Figure 7.**
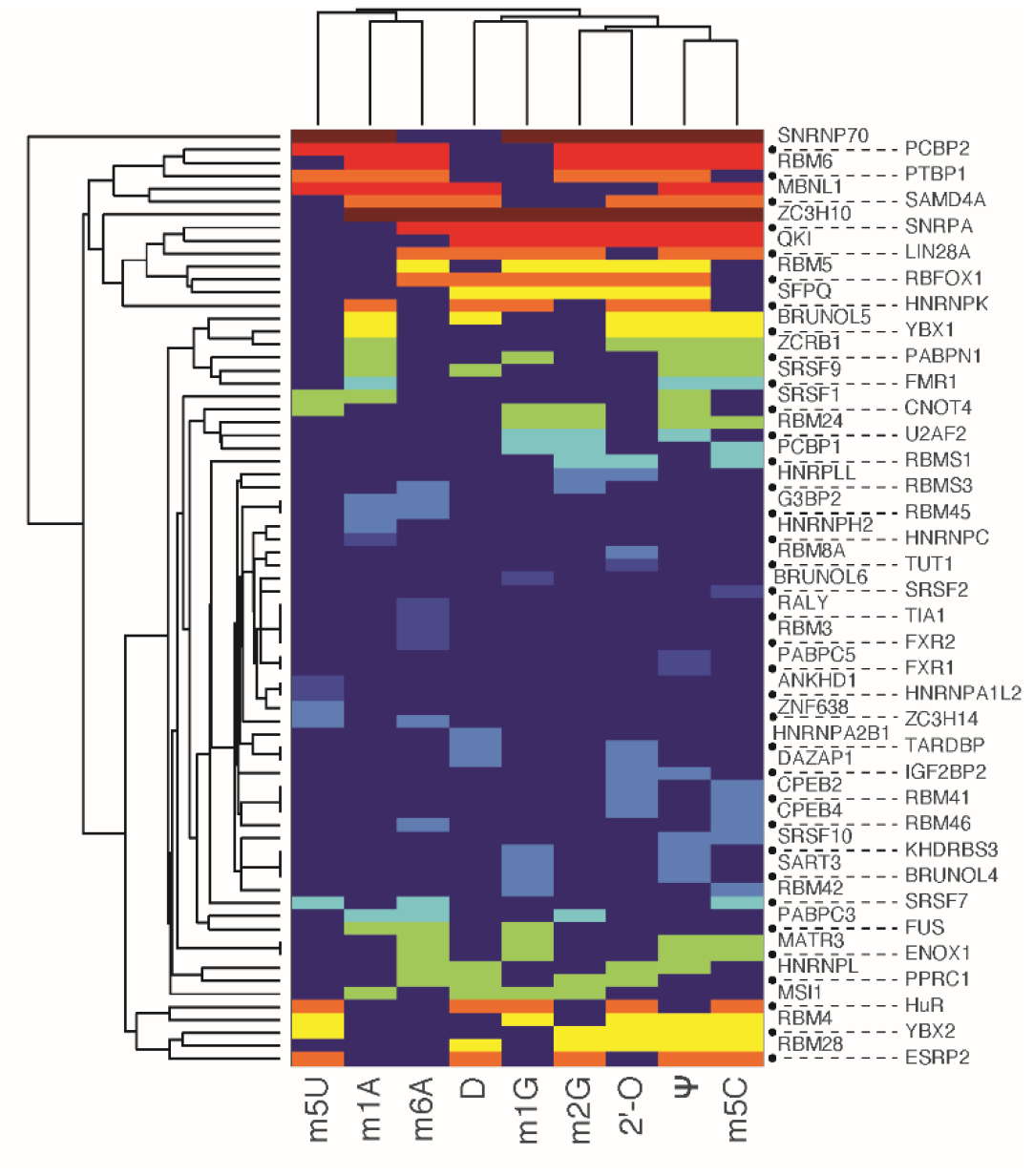
Clustergram analysis (heatmap) of RNA binding proteins identified to most likely interact with different transcriptome modifications. The color represents the significance with which they are found to be unique to these mRNA modifications by MEME-ChIP. The significance goes from lowest to highest as color varies from blue to red.

In order to further validate the credibility of the features captured by our method, we converted the 32 convolutional kernels for m^5^C samples into position frequency matrices or motifs following the similar procedure of DeepBind. Then, we aligned these motifs to known motifs using the TOMTOM algorithm [49]. Of the 32 motifs learned by the first layer of the encoder network, 25 significantly matched known RBP motifs (E < 0.05). Subsequently, we proceeded to verify whether the RBPs identified by this analysis concurs with the results of other studies investigating similar problems. In [50], the authors carried out an analysis to determine the relationship between m^5^C sites and RBPs using CLIP-seq data and reported the RBPs showing statistically significant enrichment of m^5^C in their binding sites compared to randomly sampled Cs. Expectedly, several of the RBPs identified by our motif analysis for m^5^C (6 of 25) were also discovered by that study, which further endorses the significance of our work (Supplementary Table 4).

## CONCLUSION

We considered the prediction of different transcriptome modifications based on RNA sequences. To address the problem of small sample size for many of the modifications, we developed a generative adversarial network model called MR-GAN, which is trained to learn low dimension embeddings of transcriptomic-wide sequences in an unsupervised manner. The learned embedding, as demonstrated through the experimental results, contain the improved representation of sequences for different modifications as it maps the RNA modifications with similar functionalities together. We have also demonstrated that the motifs learned by MR-GAN in the process of discriminating between various transcriptome modifications are biologically meaningful and conforms to the findings of some of the previous studies. It is noteworthy to mention that we analyzed only nine out of almost 100 well-known modifications. We believe there would be more interesting patterns revealed if the MR-GAN sequence features are applied to additional RNA modifications. The main advantage of MR-GAN is that the model can perform in a satisfactory accuracy even with the heavily skewed dataset without the need for employing data balancing techniques. This is a significant contribution to the bioinformatics research community because we often fail to develop a well-performing computation prediction model due to the lack of enough labeled data. We hope to extend this work by applying the embedding into more genomic sequence related classification problems.

## AVAILABILITY

MR-GAN is available in the GitHub repository (https://github.com/sirajulsalekin/MR-GAN). The training data are available for download at https://drive.google.com/open?id=1aASppi8f0jWk-iGNcMV1iP6yzz_tPj4g

## ACKNOWLEDGEMENT

This work was supported by the National Institutes of Health (R01GM113245 to YH, CTSA 1UL1RR025767-01 to YC, and K99CA248944 to YCC), Cancer Prevention and Research Institute of Texas (RP190346 to YC and YH, and RP160732 to YC), San Antonio Life Sciences Institute (SALSI Innovation Challenge Award 2016 to YH and YC, and SALSI Postdoctoral Research Fellowship 2018 to YCC), and the Fund for Innovation in Cancer Informatics (ICI Fund to YCC and YC). We thank the computational support from UTSA’s HPC cluster Shamu, operated by the Office of Information Technology.

## CONFLICT OF INTEREST

There is no conflict of interest.

## Notes

### Competing Interest Statement

The authors have declared no competing interest.

